# E-scape: Interactive visualization of single cell phylogenetics and spatio-temporal evolution in cancer

**DOI:** 10.1101/080622

**Authors:** Maia A. Smith, Cydney Nielsen, Fong Chun Chan, Andrew McPherson, Andrew Roth, Hossein Farahani, Daniel Machev, Adi Steif, Sohrab P. Shah

## Abstract

Inference of clonal dynamics and tumour evolution has fundamental importance in understanding the major clinical endpoints in cancer: development of treatment resistance, relapse and metastasis. DNA sequencing technology has made measuring clonal dynamics through mutation analysis accessible at scale, facilitating computational inference of informative patterns of interest. However, currently no tools allow for biomedical experts to meaningfully interact with the often complex and voluminous dataset to inject domain knowledge into the inference process. We developed an interactive, web-based visual analytics software suite called E-scape which supports dynamically linked, multi-faceted views of cancer evolution data. Developed using R and javascript d3.js libraries, the suite includes three tools: TimeScape and MapScape for visualizing population dynamics over time and space, respectively, and CellScape for visualizing evolution at single cell resolution. The tool suite integrates phylogenetic, clonal prevalence, mutation and imaging data to generate intuitive, dynamically linked views of data which update in real time as a function of user actions. The system supports visualization of both point mutation and copy number alterations, rendering how mutations distribute in clones in both bulk and single cell experiment data in multiple representations including phylogenies, heatmaps, growth trajectories, spatial distributions and mutation tables. E-scape is open source and is freely available to the community at large.

## Introduction

Cancer cell populations emerge through branched evolutionary processes leading to variation in fitness and differential growth patterns. Cell population dynamics fundamentally impact biological and clinical properties and motivate key questions for cancer focused biomedical investigators. Advances in high-throughput sequencing and computational methods ^12^ have recently enabled the quantification of clonal dynamics through analyses of how mutations distribute across cancer cell populations over temporal and spatial axes. This has led to an increased understanding of how cancer evolution impacts treatment resistance, cancer progression^3^ and metastasis^?^. Measuring cancer evolution with sequencing technology applied to both bulk tissues and single cells is a rapidly growing field and reflects the general acceptance that cancers must be studied as evolving dynamic systems in order to advance therapeutic strategies and effects.

Studying evolutionary properties of cancer with sequencing technology is rooted in the identification of (epi)genomic features which mark individual cell populations, establishing ancestral relationships of these populations and tracking their abundances in time ^4-6^ and/or anatomic space ^7-9^. Due to the statistical features of these datasets, computational scientists with high levels of mathematical and analytical sophistication are often required to interpret the data. Such skills often lie outside the expertise of biomedical investigators, who offer complementary and essential domain-specific knowledge. Interactive visualization provides a conduit for disease-focused biomedical investigators to engage with cancer evolution datasets and contribute domain-specific insights in the process of data analysis and inference. The benefits of automated, interactive visualization tools for clonal evolution are manifold: visualization systems facilitate hypothesis generation, provide stimulus for further experimentation, validate algorithmic approaches and/or expose the need for improvements. In this contribution we describe a suite of new web-based interactive visualization tools for investigators to interact with cancer genomics datasets focused on clonal evolution.

The nature of clonal evolution datasets varies by experimental design. An important quantitative measure is the variant allele frequency (VAF) of a mutation from NGS experiments. This typically equates to the number of reads harbouring a point mutation divided by the total number of reads aligning to the locus of interest and is accepted as a first approximation to the proportion of alleles present in the DNA substrate. When measured across several mutations, a number of approaches have been developed to convert allele prevalences to mutation cellular prevalences: defined as the proportion of cells in the input material harbouring the mutation of interest. This quantity is also referred to as cancer cell fraction (CCF) and can be computed by a number of different methods ^2^, which aim to account for confounding factors impacting allele prevalence: the proportion of non-malignant cells in the sample, concomitant copy number alterations and expectation that subsets of mutations will share the same cellular prevalence.

Typical assumptions state that each mutation is harboured by a population of cells which share the same underlying genotype (to varying degrees of precision). When mutation prevalence is measured across multiple tumour samples, groups of mutations may vary consistently in prevalence, yielding powerful signals to identify genotypes. Quantifying genotypes in turn, marks constituent clones in the samples and their clonal prevalence changes over the key dimensions of time and space. It naturally follows that measurements of temporal samples, such as serial patient biopsies or serially propagated patient derived xenografts, reflect clonal growth dynamics, and measurements over spatial samples (metastases and physically separated anatomic samples via multi-region sampling) reflect distributions of clones as a function of tumour microenvironments. Phylogenetic associations of clones can also be inferred from multiple sample experimental designs thereby establishing evolutionary histories and ancestral relationships of clones.

Clonal measurements are also obtained via single cell sequencing. Single cell measurements can be executed across the whole genome of a cell or through targeted measurement of a specific set of loci. Whole genome assays typically yield copy number profiles, ascertaining segmental aneuploidies while targeted assays yield presence/absence (or locus specific genotype) data on per cell and per targeted mutation locus dimensions. These data can be input into algorithms for single cell phylogenies relating the evolutionary relationships between cells in the sample of interest.

Together, clonal prevalences over a set of mutations (single or multiple samples over time or space), clone and single cell phylogenies and cell-by-locus matrices form a multi-faceted set of inputs for biological interpretation. Although the field is partially served by methods for visualizing clonal expansions ^10^ over time and for single cell copy number profiles ^11^, no interactive data visualization software currently exists to allow the biomedical investigator to interactively navigate these datasets, nor the various facets.

Most often, static plots of data - including dendrograms coupled with cell-by-locus heat maps, representations of mutations in specific cells and various depictions of clonal compositions across space and time - are rendered for publication or presentation. This introduces a challenge - various figures and tables contain linked information but in a static visualization regime, obvious barriers to interactivity limit the ability to exploit these links for enhanced interpretation. Interactive visualization introduces browseability of data. For example, selecting part of a clone phylogeny could identify where in anatomic space of a metastatic patient that clade is represented; selecting part of a cell-by-locus heat map to determine the phylogenetic topology of a group of cells; or interactively determine which mutations associate with an expanding clone over time. Linked, dynamically updating plots capable of responding to user-driven actions are required to support this nature of data interactivity.

To fill this need, we developed the E-scape (Evolutionary Landscapes) tool suite. E-scape consists of three automated visualization tools: i) TimeScape for longitudinal, timeseries analysis; ii) MapScape for spatial distribution analysis of metastases; and iii) CellScape for single cell analysis including phylogenetic and population dynamics of cancer. The suite of tools allows, for the first time, interactive visualization and browsing of clonal evolution datasets. The tools are freely accessible, and are easily integrated into web documents for sharing. Launched from R as htmlwidgets^12^, the output web-based visualizations support interactivity with the data through brushing and linking operations in a web browser, facilitating fully integrated views of multiple data representations of clonal evolution studies.

## Results

### Interactive visualization software suite for cancer evolution

We developed a novel software platform for visualising clonal evolution datasets, schematically represented in Figure S1. The platform is engineered with two main components: a data handling component developed in R^13^ (version 3.2.2), and a web component written in JavaScript (version 1.7). Using an R-based platform facilities integration with other popular packages in the bioinformatics community, while JavaScript provides a route to powerful web based graphing libraries, specifically d3.js^14^ (version 3.5.6). An interface between the two components, the R htmlwidgets framework^12^ (version 0.6), produces html output (optimized for Chrome version 52.0.2743) from user input in R.

The internal design of each E-scape tool consists of a consistent layered architecture. In the data handling component, the R htmlwidget receives inputs in the form of data tables and strings. Tabular data representing mutations are then checked for integrity and converted into pixel matrices for rendering. The data are then transformed into a list of JavaScript Object Notation (JSON) elements in preparation for migration to the web component. The web component transforms the data into d3.js-compatible formats, which performs two functions: (i) mapping data elements to their final visual positions and forms and (ii) defining user interactivity. The resultant html interface for data exploration appears in the user’s browser (see Online Methods for visualization interactivity and sharing).

Each E-scape interface contains a similar layout (e.g. Fig. 1). A toolbar housing interactive tools sits atop each page. The main view contains a dashboard of linked visualizations of the data. A legend appears on the right-hand side of the dashboard, and is linked to the main view to enable highlighting of categorical elements such as clones or sampling time points.

**Figure 1:**
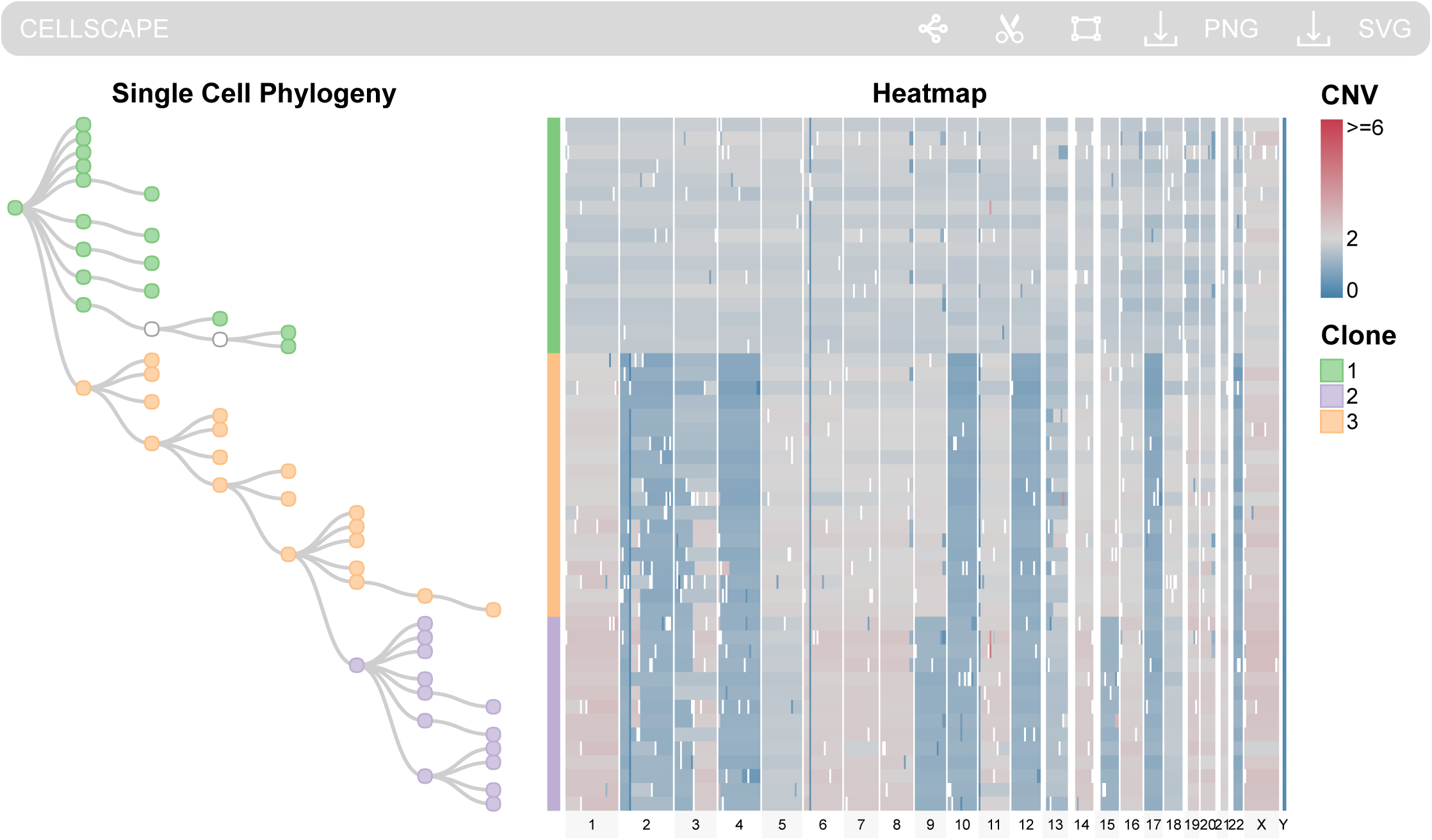
A CellScape visualization of triple-negative breast cancer single cell data from Wang *et al*.^15^. Supplementary Link 11 shows an interactive version of this view.

Within each E-scape tool, we implemented a number of interactive features to expose the additional relationships that may not be immediately apparent from a static 2D representation. These features also empower the user with additional quantitative details, multiple perspectives, filtering and editing capabilities.

### CellScape: Visualizing clonal evolution at single cell resolution

CellScape facilitates interactive browsing of single cell clonal evolution datasets. The tool requires two main inputs: (i) the genomic content of each single cell in the form of either copy number segments (see Supplementary Table 1) or targeted mutation values (see Supplementary Table 2), and (ii) a single cell phylogeny (see Supplementary Table 3). Phylogenetic formats can vary from dendrogram-like phylogenies with leaf nodes to evolutionary model-derived phylogenies with observed or latent internal nodes. The CellScape phylogeny is flexibly input as a table of source-target edges to support arbitrary representations, where each node may or may not have associated genomic data. The output of CellScape is an interactive interface displaying a single cell phylogeny and a cell-by-locus genomic heatmap (Fig. 1) representing the mutation status in each cell for each locus. These two views complement one another: the phylogeny expresses the evolutionary progression from one clonal lineage to another, and the heatmap displays the cell-specific large- and small-scale genetic changes as cell populations evolve. The default view visually links the phylogeny with the heatmap by vertically aligning the phylogenetic nodes to their genomic profiles (Fig. 1).

To exemplify the CellScape view, we imported triple-negative breast cancer single cell data from Wang *et al.*^15^ (Fig. 1). This visualization depicts an evolutionary progression from the mostly diploid green ancestral clone (presumed non-malignant cells) to the orange clone with acquired genomic losses in chromosomes 2, 3, 4, 10, 12, 17 and 22, and finally to the purple clone with additional losses and gains throughout the genome.

Phylogenetic tree representations are diverse and depend on the algorithm which generated them. We therefore implemented a flexible suite of phylogenetic views supporting varying outputs that can be anticipated from the range of algorithms (Fig. S2). The force-directed graph view (Fig. S2c) displays an unrooted phylogeny. Alternatively, the unidirectional tree view (Fig. S2a) displays a rooted phylogeny and vertically aligns nodes to their corresponding cells in the heatmap (Fig. 1). The phylogenetic views also support edge distance information to emphasize evolutionary distances encoded by branch lengths (Figs. S2b and S2d). In CellScape’s phylogenetic view, nodes are interactively linked to their genomic profiles. For *individual cells,* we achieve this by mouseover of any element. For *groups of cells,* we created a selection tool to select genomic profiles of interest and highlight the corresponding nodes in the phylogeny (Fig. S3). For *clones,* whose cell populations may be intermixed in the phylogeny, we designed the legend such that mouseover of any clone will reveal its presence in the phylogeny and heatmap.

**Figure 2:**
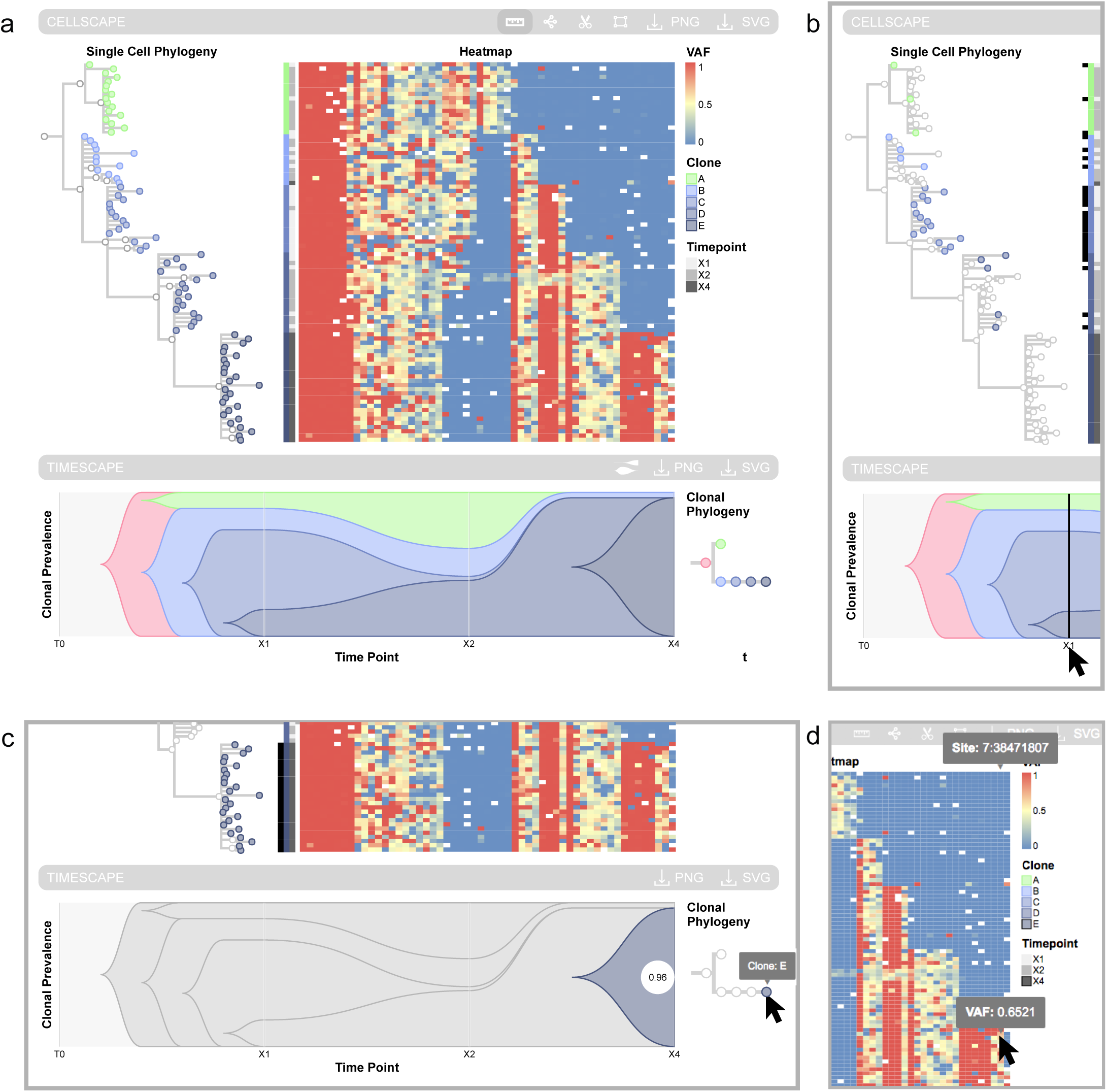
A time-series CellScape visualizes the xenograft SA501 single cell data published in Eirew *et al.*^4^ For the interactive view, see Supplementary Link 1. (a) The full time-series CellScape view. The bottom panel displays a TimeScape automatically generated from the single cell data. (b) Mouseover of time point “X1” elicits a response in “X1”-associated nodes and heatmap rows, emphasizing the diversity of cells at this time point. (c) Mouseover of a clone causes reactive highlighting in the TimeScape, single cell phylogeny, and heatmap. (d) The user inspects a heatmap cell for its Variant Allele Frequency (VAF), mutation site and single cell ID (not shown).

A major challenge for single cell data is missing or low-quality data. We implemented a tree trimming tool (Fig. S4) which can be used to interactively remove unwanted branches of the phylogeny and their corresponding genomic profiles. This is especially useful for clades that may be grouped on account of noise in the measurement rather than true biologically relevant variation.

Within the phylogeny, cells from different time points may be intermixed such that it is difficult to interpret how the tumour evolved over time. We designed CellScape to accommodate temporal sampling information by automatically generating and appending a TimeScape to the view. The TimeScape provides a context of temporal clonal dynamics while the CellScape presents a granular view of clonal and temporal constituents. To illustrate this, we imported the single cell data from serially propagated patient derived breast cancer xenografts^4^. We depict SA501 in CellScape (Fig. 2), showing how on intitial engraftment, the tumour cell population is diverse with multiple co-existing clones (Fig. 2b). However, later passages reveal that a uniform cellular population (Fig. 2c) which was rare at early timepoints, has undergone a selective sweep, thereby constituting a dominant population (Fig. 2d). The CellScape view therefore at once can link single cells from one or several serially sampled timepoints, giving insight into cell populations dynamics within the context of a phylogeny.

### TimeScape: visualization of temporal clonal evolution

TimeScape (Fig. 3) is an automated tool for navigating temporal clonal evolution data. The key attributes of this implementation involve the enumeration of clones, their evolutionary relationships and their shifting dynamics over time. TimeScape requires two inputs: (i) the clonal phylogeny (see Supplementary Table 4) and (ii) the clonal prevalences (see Supplementary Table 5). Optionally, TimeScape accepts a data table of targeted mutations observed in each clone and their allele prevalences over time (see Supplementary Table 6). The output is the TimeScape plot showing clonal prevalence vertically, time horizontally, and the plot height optionally encoding tumour volume during tumour-shrinking events (Fig. 3a). At each sampling time point (denoted by a faint white line), the height of each clone accurately reflects its proportionate prevalence. These prevalences form the anchors for bezier curves that visually represent the dynamic transitions between time points. We note that bezier curves are simple approximations for the underlying sparsely-sampled biology, and may not accurately reflect the dynamics between samplings. Future research modeling patterns of clonal dynamics with more granular sampling intervals will better approximate growth kinetics.

**Figure 3:**
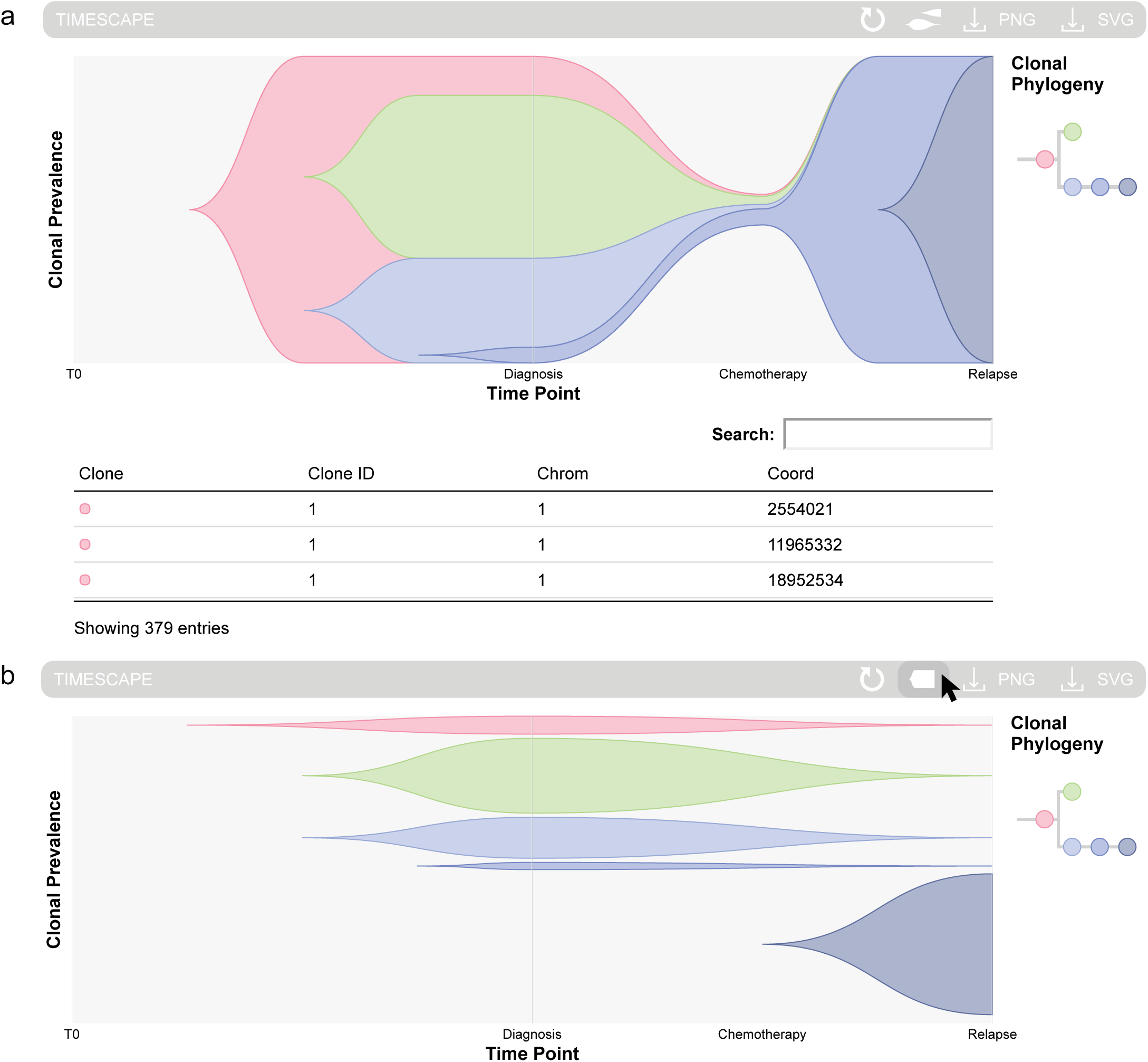
A TimeScape displaying the clonal evolution as expressed in AML patient 933124 from Ding *et al.*^11^ For the interactive TimeScape, see Supplementary Link 8. (a) In the nested view, descendant clones emerge from their ancestors. (b) In the clonal trajectory view, each clone changes in prevalence on its own track. The clonal hierarchy is maintained by the vertical ordering and horizontal offsetting of clones.

We designed TimeScape to include two additional outputs, the clonal phylogeny and (if provided) the mutation table, to provide evolutionary and genomic context, respectively. In order to correlate mutational content, clonal prevalence and the phylogenetic relationships between clones, we implemented interactive linking of the three views in TimeScape. Clicking clones in the phylogeny will reveal their specific mutational content in the mutation table (Fig. S5a), and clicking each mutation will reveal its variant allele frequency across time (Fig. S5b). The plot is designed to preserve clonal hierarchy by nesting clones according to their branching topology (Fig. 3a). However, the nested nature of clones can confuse the perception of clonal prevalence between sampling time points. For instance, when a descendant clone replaces its ancestor, there is a false illusion of ancestral expansion (Fig. 3a, mid-blue clone). To clarify the trajectories of individual clones, we implemented two interactive features in TimeScape: (i) the user may mouseover each clone to display its temporally varying cellular prevalence (Fig. 2c), and (ii) the nested organization of clones may be interactively disabled, revealing a new perspective that assigns each clone to a separate track while retaining hierarchical intuition through intelligent vertical ordering and horizontal offsetting of clones (Fig. 3b).

The automated nature of TimeScape offers great potential for studies with large sample sizes. By easily and quickly plotting TimeScapes of many patients with the same condition, patterns of clonal dynamics can emerge. For example, Kridel *et al.*^3^ have analyzed the clonal dynamics of follicular lymphoma patients that have undergone a histological transformation to aggressive disease. Their results show that the majority of transformed follicular lymphomas exhibit dramatic clonal dynamics upon histological transformation. TimeScape clearly displays this finding by quickly plotting the fifteen patients in their study, revealing that post-transformation dominant clones originated as minor clones prior to transformation (Fig. 4), and in most cases, clones constituting the original diagnostic specimens are from clades on opposite branches of the clonal phylogeny. Automation of TimeScape is also valuable for many serially sampled specimens (e.g. xenograft studies where serial transplantations provide longitudinal tracking of clonal dynamics in a cancer, or serial biopsies from patients). Figure 2 and Supplementary Link 1 both illustrate a TimeScape of the xenograft single cell data published in Eirew *et al.*^4^ consisting of three timepoints, although plotting longer timeseries with more granular intervals is equally supported (Fig. S6).

**Figure 4:**
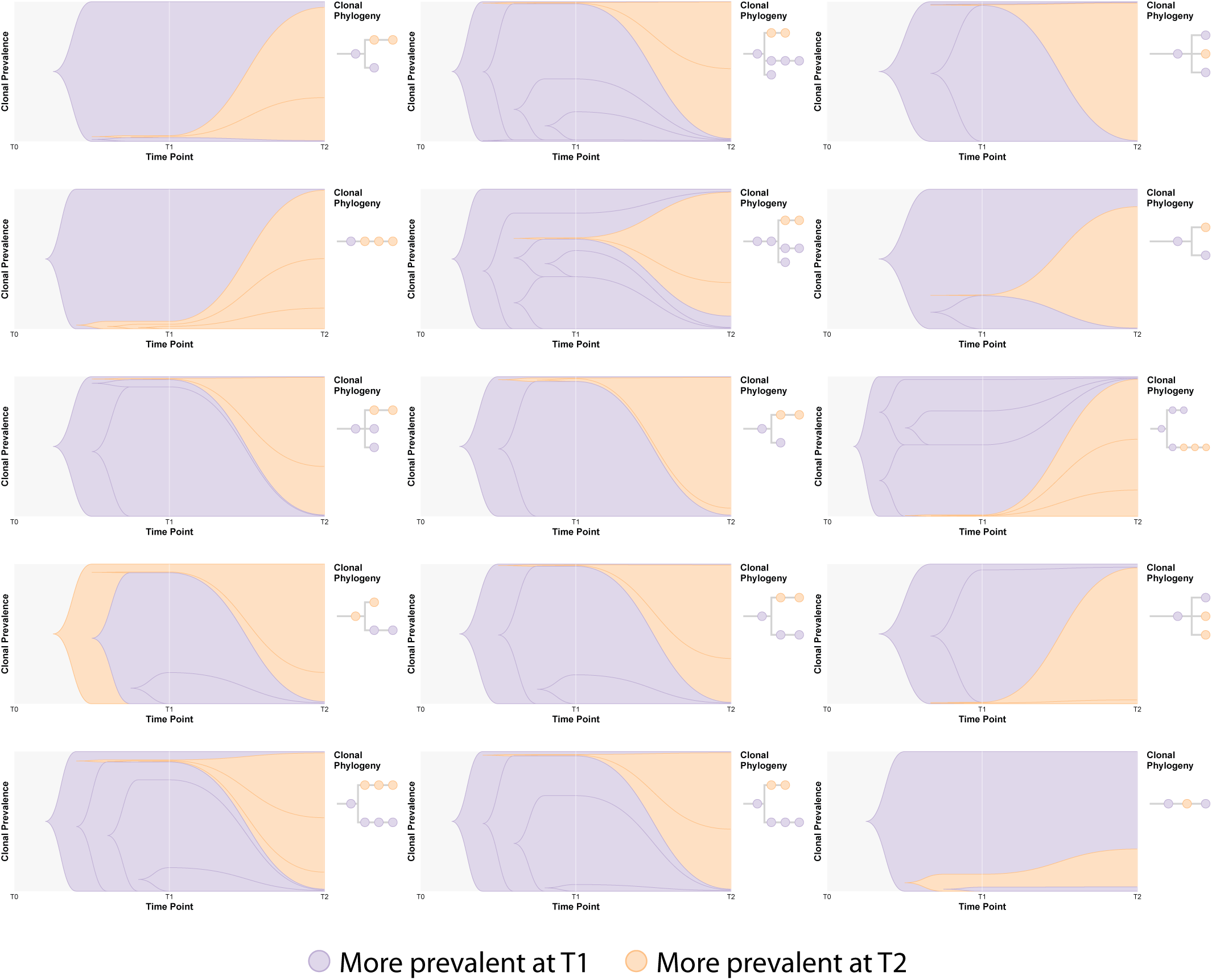
TimeScapes of transformed follicular lymphoma patients from Kridel *et al.*^3^ The clones more prevalent at time point 1 are shown in purple, while those more prevalent in time point 2 are shown in orange. From time point 1 to time point 2, there is a clear shift in clonal composition in fourteen out of fifteen patients, where minor clones at time point 1 come to dominate at time point 2. For an interactive version of these TimeScapes, see Supplementary Link 12.

### MapScape: Visualization of spatial clonal evolution

MapScape integrates clonal prevalence, clonal hierarchy, anatomic and mutational information to provide interactive visualization of spatial clonal evolution. A strong interest in spatial series has emerged in the literature, owing to capacity to identify mutations constituting the root of phylogenetic trees and those that are present in subsets of cells ^16^; and reconstructing clonal migration patterns of pre-treated ^7^ or autopsy specimens ^8^. There are four inputs to MapScape: (i) the clonal phylogeny (see Supplementary Table 7), (ii) clonal prevalences (see Supplementary Table 8), (iii) an image reference, which may be a medical image (see http://mo_bccrc.bitbucket.org/escape/images/tumours_A21.png) or drawing (see http://mo_bccrc.bitbucket.org/escape/images/patient_1.png) and (iv) pixel locations for each sample on the referenced image (see Supplementary Table 9). Optionally, MapScape can accept a data table of mutations for each clone and their variant allele frequencies in each sample (see Supplementary Table 10). The output of MapScape (Fig. 5) consists of a cropped anatomical image (see Online Methods Section “Image auto-crop in MapScape”) surrounded by two representations of each tumour sample. The first, a cellular aggregate inspired by Eirew *et al.*^4^, visually displays the prevalence of each clone. The second shows a skeleton of the patient’s clonal phylogeny while highlighting only those clones present in the sample. Together, these representations enable the analyst to visualize the distribution of clones throughout anatomic space.

**Figure 5:**
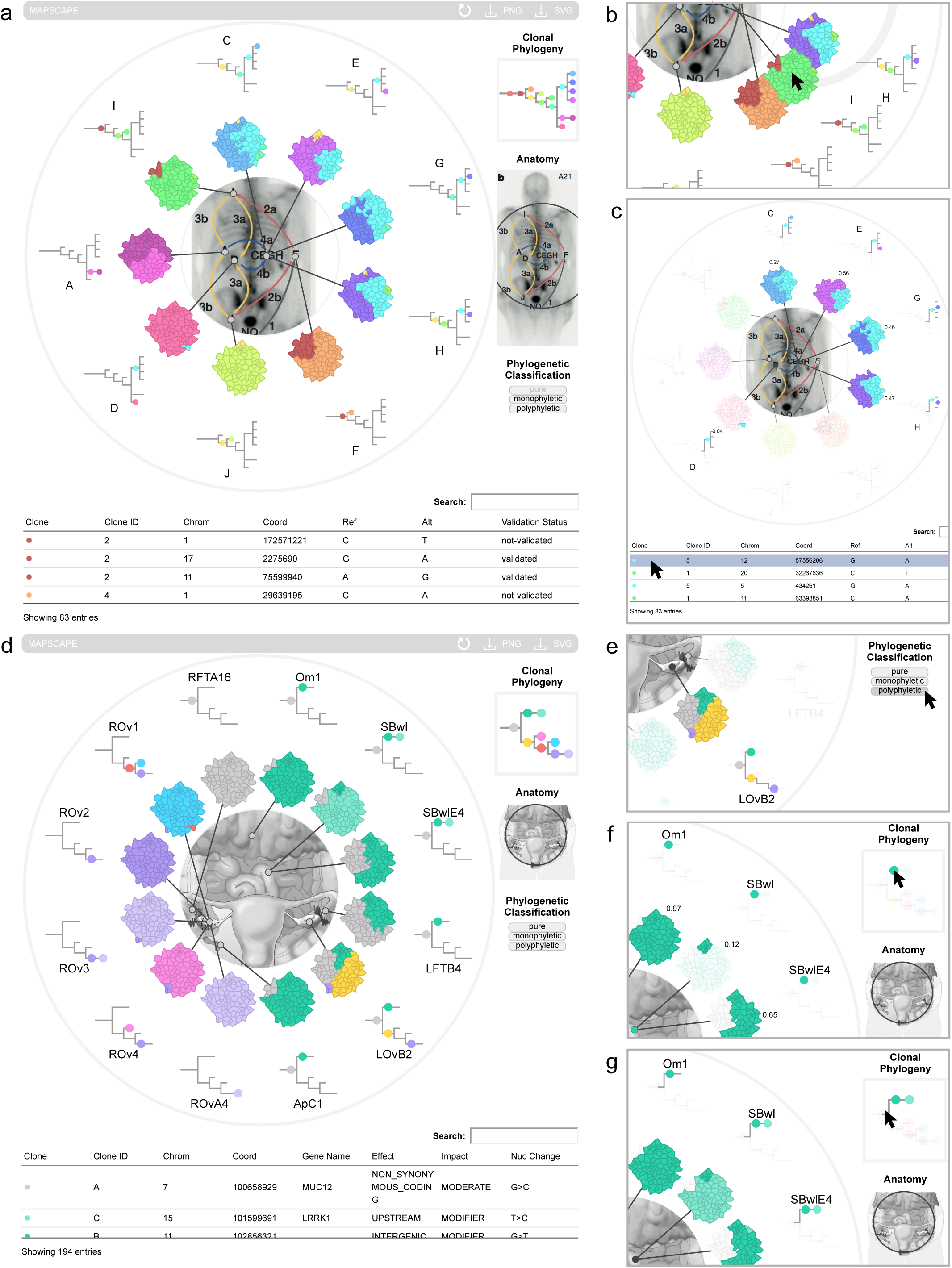
An overview of MapScape. A MapScape visualization of metastatic prostate cancer data published in Gundem et *al.*^8^. For an interactive version of this view, see Supplementary Link 3. For sample reordering purposes, the user may click and drag a sample. This feature may be used to emphasize effects such as metastatic progression between samples. Clicking a mutation displays its sample-specific variant allele frequencies at each participating tumour sample. A MapScape visualization of metastatic ovarian cancer Patient 1 published in McPherson et *al.*^7^ For an interactive version of this view, see Supplementary Link 4. Mouseover of a phylogenetic classification label highlights all tumour samples following this classification. Mouseover of a clone highlights the clone throughout the view and displays its prevalence in each sample. Mouseover of a phylogenetic branch highlights the descendant clones throughout the view.

Surrounding the main view, the interface includes evolutionary, mutational and anatomical contexts via the clonal phylogeny, the mutation table and the full anatomical image, respectively. These contexts relate to one another facilitating dynamic, interactive linking and browsing between all elements. Interacting with the clonal phylogeny enables a clone- or lineage-centric view, where the user may identify each clone’s spatial distribution and prevalence in the main view, as well as its specific mutational content in the mutation table. As a concrete example, mouseover of the green coloured clone in Patient 1 from McPherson *et al.*^7^ highlights the distribution of this clone within the patient (Fig. 5f), and mouseover if its preceding branch displays the complete descendant lineage (Fig. 5g). Interacting with the mutation table exposes the phylogenetic and anatomic distribution and emergence of mutations. For example, selecting a mutation in Patient A21 from Gundem *et al.*^8^ displays the clones and samples harbouring the mutation, from which users could infer its anatomical origin (Fig. 5c, site D) and potential influence on tumour progression (Fig. 5c, blue and purple clones).

We implemented a default sample layout method (see Online Methods, Section “Auto-layout of samples in MapScape”), however, we allow the user to radially reorder samples in order to clarify certain effects. Figure 5b shows the user reordering samples for a logical metastatic progression. We enable user interaction with each of the legend’s phylogenetic classification labels in order to highlight the local structure of each tumour sample. For instance, mouseover of the “Polyphyletic” classification highlights the left ovary, a site of high clonal diversity that exhibits two phylogenetic branches emanating from the ancestral clone (Fig. 5e).

Within a MapScape visualization, clicking multiple clones in the clonal phylogeny will highlight their presence throughout the view. This multi-clone selection feature empowers the user to visually group clones with similar properties. Figure S7 and Supplementary Link 2 demonstrate the use of this feature to map the clonal distribution and progression of Patient 3 from McPherson *et al.*^7^, a paper where MapScape was used to visually summarize each metastatic cancer patient. Within this patient, a subset of clones are present in all eight sampled locations—these clones developed early and possessed high metastatic potential (Fig. S7a). Later in development, descendant clones emerged and migrated to one or more sites in the patient (Fig. S7b). The latest clones in development are private to individual anatomical locations, their confinement due to a lack of either metastatic potential or time (Fig. S7c).

MapScape can be used to analyze metastatic progression from one site to another within a patient. As a biological vignette, we imported two datasets displaying varying modes of tumour cell seeding. The first dataset describes metastatic prostate cancer Patient A21 from Gundem *et al.*^8^. The MapScape of this data reveals polyclonal seeding—sample H likely received the green clone from sample I, the blue clone from sample D, and the yellow clone from sample J (Fig. 5a-c, Supplementary Link 3). The second dataset describes metastatic ovarian cancer Patient 1 from McPherson *et al.*^7^ (Fig. 5d-g, Supplementary Link 4). Analysis of this MapScape suggests that all sites were seeded from the left ovary except the left ovary itself, which was re-seeded by the right ovary. A possible progression sequence is as follows: the yellow clone migrated from the left ovary to the right ovary, this clone was then replaced by its descendants, and one of these descendants—the purple clone—then re-seeded the left ovary.

When a dataset contains both spatial and temporal components, MapScape and TimeScape can combine to form an additional interpretation layer. As an illustration, we imported data from Patient 7 published in McPherson *et al.*^7^ and display both spatial and temporal visualizations in Figure 6. This figure shows that the green clone, despite its minority (Fig. 6b, time point “Intraperitoneal diagnosis”), spread to the brain via hematogenous dissemination from the right uterus (Fig. 6a, site “RUtD3” and brain sites). The figure also makes evident that surgical interventions reduced but did not eliminate the clonal diversity in the intraperitoneal space (Fig. 6b, time points “Intraperitoneal diagnosis” and “Intraperitoneal relapse”). The relapsed clones—rather than recolonizing resected sites—colonized new areas of the intraperitoneal space (Fig. 6a, sites “RPvM” and “BwlA6”).

**Figure 6:**
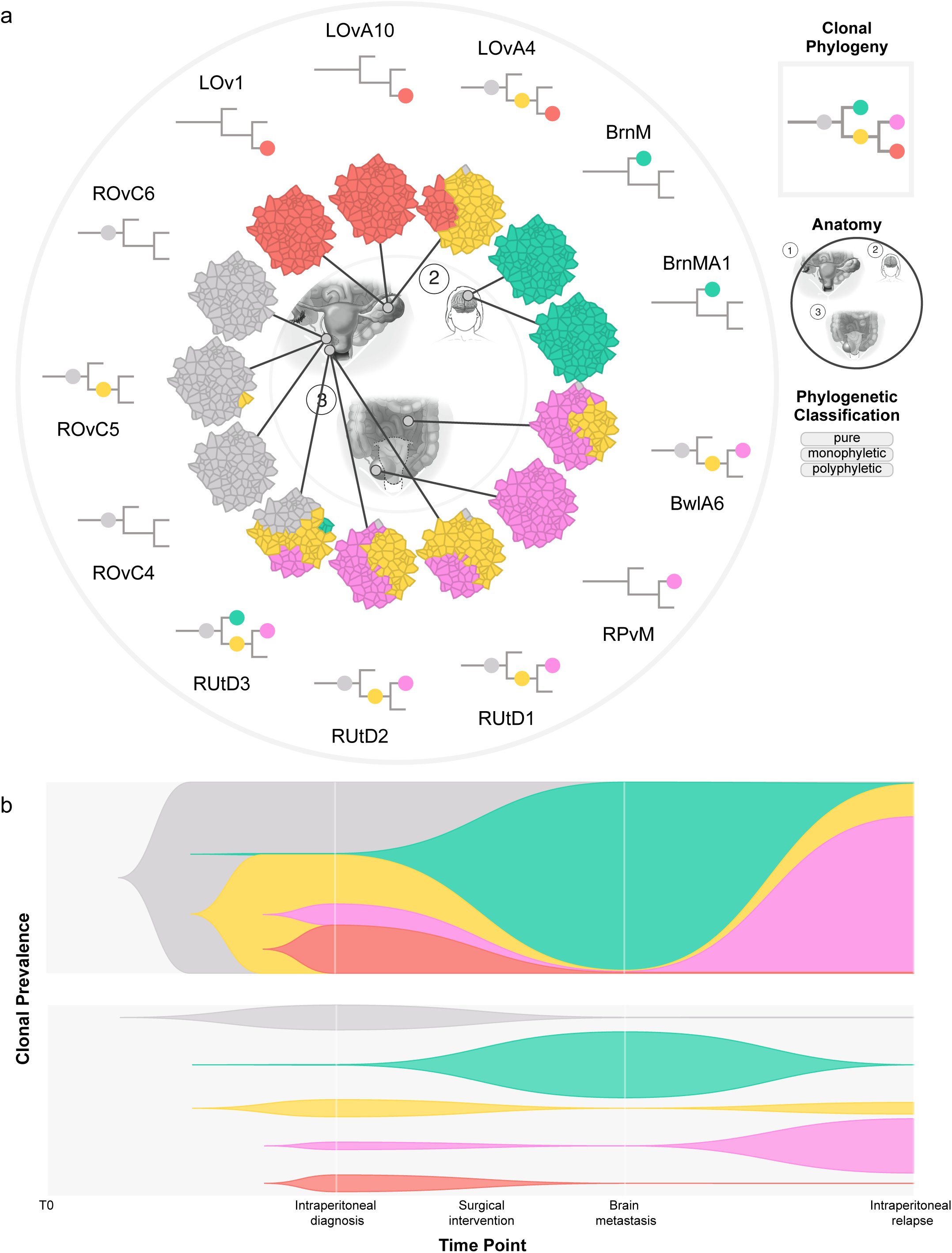
Both time-series and spatial visualizations of metastatic ovarian cancer patient 7 from McPherson *et al.*^7^. Together, these views highlight two different avenues of metastatic progression in the patient: intraperitoneal spread and hematogenous dissemination. (a) A MapScape visualization displays the samples taken from a variety of metastases over three time points. The view reveals the metastatic potential of the right uterus tumour, as it colonizes the brain via hematogenous dissemination, and also colonizes the bowel and right pelvic mass via intraperitoneal spread. Interactive MapScape available at Supplementary Link 13. (b) A TimeScape visualization shows the clonal composition at each sampling time point, highlighting the dramatic global temporal dynamics in this patient. At “intraperitoneal diagnosis”, a diversity of clones inhabit the peritoneum. After surgical interventions, the pink and yellow clones survive; at “intraperitoneal relapse” these clones seed and come to dominate the right pelvic mass and bowel implant. Likely hematogenous dissemination of the green clone from the right uterus results in a clonally-pure “Brain metastasis”. Interactive TimeScape available at Supplementary Link 14.

### Runtime Performance

The runtime performance of TimeScape and MapScape is trivial - these datasets are never large. Single cell datasets, on the other hand, typically consist of hundreds of cells, a scale that CellScape can efficiently process. For instance, visualizing our largest copy number dataset of 371 single cells and 37,580 copy number segments took 5.433 seconds of CPU time on a 2.8 GHz Intel Core i7 Macintosh computer, and less than 5 seconds to load on a Chrome browser (version 52.0.2743). Targeted mutation datasets are more manageable, as there will typically be fewer targets than copy number changes per cell. For instance, the SA501 xenograft data from Eirew *et al.*^4^ comprises 106 single cells each with only 55 mutations, and takes 0.373 seconds of CPU time on the same machine, and less than one second to load on the same Chrome browser. Thus, for currently available datasets, CellScape does not require external machinery, and any biomedical researcher can operate the tool on their personal computer.

## Discussion

The nature of cancer evolution data suggests that biological interpretation will be improved by the involvement of biomedical experts in the inference and analysis process. However,due to scale and complexity of the data, this remains intractable in the absence of automated and interactive visualization tools. We developed the E-scale tool suite: the first interactive and automated set of visualization tools for the exploration of clonal dynamics in cancer. We anticipate E-scape will aid the discovery process through visualization.

The novel contributions provided by E-scape are manifold. Beyond its core ability to visually represent clonal evolution data, E-scape interactively (i) links cell population-level clonal dynamics with a single cell resolution perspective, (ii) visualizes temporal and spatial clonal dynamics of relapsed and metastatic cancer samples, and (iii) improves accessibility of clonal evolution datasets to all researchers. We expect that with these contributions, E-scape will provide the impetus for novel insights into biological properties of evolving cancer cell populations from patients at the levels of academia, the clinic, commercial entities and perhaps even the general public.

The current release of E-scape has several limitations. Future improvements include incorporation of a backend indexing platform for improved scalability. As the magnitude of experimental output increases to tens of samples, thousands of mutations and millions of single cells, we expect that millisecond responses in browsers will be required with dynamic queries generated by user action. This will allow the E-scape to render data of arbitrary size and volume. The integration of E-scape with other plot types is another priority for facilitating alternative granular perspectives on the data. For example, an integrated whole genome browser would provide a more detailed view of clonal and single cell genotypes, sharpening copy number breakpoints and revealing small copy number changes. The incorporation of data quality metrics would facilitate the separation of signal from noise. Visualization support for emerging data types, such as single cell expression data and single cell *in situ* PCR data, will become critical as their availability becomes increasingly common in the field.

We anticipate many applications for E-scape in a variety of settings. Within research labs, E-scape is effective for data analysis and interpretation, as is thoroughly explored in this contribution. In the broader scientific community, E-scape is valuable for communicating research findings, given its capability to produce publication-ready figures that straightforwardly present complex research and straightforward deployment to the web. In the clinic, E-scape visualizations may aid clinical decision-making in the future of personalized oncology. Visualizing dynamic clonal composition at diagnosis and relapse may mechanistically explain disease progression and/or influence how oncologists intervene with alternate treatment strategies. Several recent literature findings indicate a strong rationale for ascertaining the clonal dynamics of disease during the clinical course. Clinicians will need tools such as E-scape to implement this in practice.

E-scape helps to facilitate a multidisciplinary analysis process whereby computational biologists, biomedical researchers and clinicians can effectively engage with genomic datasets through intelligent interactive and automated visualization. We view E-scape as a vehicle to bring together all components of studying dynamics of disease progression ultimately leading to biological understanding and clinical benefit.

## Methods

### Execute E-scape visualizations in this paper

Scripts to execute the E-scape visualizations appearing in this paper are available on github (see Online Methods Section “Code availability”). The scripts are visible within the documentation of each E-scape tool, and each visualization may be created using the *example* function (see Online Methods Section “Runtime execution of E-scape tools”).

## Data for prostate cancer Patient A21 (Gundem *et al.*^8^)

### Learning the clonal prevalences

In their paper, Gundem *et al.* group the clonal and subclonal mutations by their cancer cell fractions (CCFs), the fraction of tumour cells carrying a specified mutation, in each sample. For each patient, CCFs from sample pairs are plotted against one another to determine the clonal phylogeny for the patient. For each sample, we combine these two sources of information—the sample-specific CCFs and the patient clonal phylogeny—to infer clonal prevalences in a process summarized by Figure S8. Similarly to their paper, we assume the infinite sites hypothesis, which postulates an infinite number of mutation sites, although a mutation may only occur once at any given site and cannot revert to its previous form. From this assumption follows the additivity assumption, where mutations in an ancestral branch will be present throughout their descendant lineages while novel mutations accumulate. Therefore, to obtain the prevalence of each clone, we sum the CCFs of mutation clusters originating in the immediately descendant branches, then subtract this sum from the CCF of the mutation cluster originating in the immediately ancestral branch. The resulting clonal prevalences are available in the Supplementary Table 8.

### Mutational information

The mutation information was taken from their paper in the Supplementary Information section, Supplementary Data, tab “subs”.

### Clonal phylogeny

The clonal phylogeny was taken directly from Figure 2 of their paper (see Supplementary Table 7).

### Medical image and sample locations

The medical image of Patient A21 was cropped from Figure 3 of their paper, and is available at Supplementary Link 5. The locations of each sample are noted in Figure 3 of their paper, and the pixel coordinates for each location are available in Supplementary Table 9.

## Data for ovarian cancer Patient 1 and Patient 7 (McPherson *et al.*^7^)

### Clonal prevalences

Clonal prevalences were taken from Supplementary Table 9 of their paper.

### Patient 7 time-series clonal prevalences

To calculate time-series clonal prevalences for TimeScape, we averaged the prevalence of each clone (reported in Supplementary Table 9 of their paper) over all samples for each time point. The resulting clonal prevalences are available in Supplementary Table 11.

### Clonal phylogeny

The clonal phylogenies were reported in Figure 4 of their paper, and are available in Supplementary Tables 12 and 13.

### Mutations, their clonal assignments and their prevalences

Mutations, their clonal assignments and prevalences in each sample were obtained from the authors of the paper, and are available in Supplementary Tables 14 and 10.

### Medical images and sample locations

The anatomical diagrams of Patients 1 and 7 were obtained from the authors of the paper, and are available at Supplementary Link 6 and Supplementary Link 7. The locations of each sample are noted in Figure 4 of their paper, and the pixel coordinates for each location are available in Supplementary Tables 15 and 16.

#### Data for acute myeloid leukemia Patient UPN 933124 (Ding et *et al*.^17^)

The clonal prevalences and phylogeny of Patient UPN 933124 were both reported in the main text of the paper, and are available in the Supplementary Tables 5 and 4, respectively. The deep readcounts of somatic mutations were taken from Supplementary Table 5a in their paper (VAFs were taken from the “avg.tum.var.freq” column), and are available in Supplementary Table 6.

#### Data for transformed follicular lymphoma patients (Kridel et *al.*^3^)

The clonal prevalences and phylogeny of the transformed follicular lymphoma patients were obtained directly from the author, and are available in Supplementary Tables 17 and 18.

#### Data for xenograft SA501 (Eirew et *al.*^4^)

The targeted mutations and single cell phylogeny were obtained directly from the authors of the paper, and are available in Supplementary Tables 2 and 3, respectively.

## Data for triple-negative breast cancer patient (Wang et *al*.^15^)

### Copy number data

The single cell copy number data was obtained directly from the authors, and is available in Supplementary Table 1.

### Learning the phylogenetic tree

We learned the phylogenetic tree from the single cell copy number data. In traditional phylogenetic algorithms, e.g. neighbour-joining, the internal nodes and leaves of the tree are assigned to unobserved ancestors and observed genotypes, respectively. However, we used a novel algorithm that can assign observed genotypes to internal nodes. This is an important capability for single cell cancer data, where both descendants and ancestors can be observed in a sample. The phylogeny is available in Supplementary Table 19.

#### Code availability

TimeScape, MapScape and CellScape are open source and available on github at https://bitbucket.org/MO_BCCRC/timescape, https://bitbucket.org/MO.BCCRC/mapscape and https://bitbucket.org/MO_BCCRC/cellscape, respectively.

#### Runtime execution of E-scape tools

To run the E-scape tools, the user must have R (version >= 3.2.2) installed on their computer. The user then runs the following commands:

~~~
                               **install.packages**(”devtools”)
                               **library**(devtools)
                               **install_bitbucket** (”MO_BCCRC/<tool_name>”)
                               **library**(<tool_name>)
~~~

To view the input requirements (Fig. S1, Supplementary Tables 20, 21 and 22) and example code for each tool, the user runs the following command:

~~~
                               ?<tool_name>
~~~

To run the example code for each E-scape tool, the user types the following command:

~~~
                               **example**(<tool_name>)
~~~

#### Interacting with the visualizations

The main view in MapScape displays a patient's tumour samples centrally connected to an anatomical diagram. The content and interplay of these samples may be interactively explored by the user. Mouseover of an anatomical location highlights its associated tumour samples and displays its location title (e.g. Right Ovary). Furthermore, dragging the tumour samples radially will enable the user to emphasize effects such as the progression of metastasis.

The main view in a TimeScape is capable of displaying two different representations of a tumour's clonal composition changing over time, but only displays one representation at a time. A toggle is available for switching between these two representations.

The legends for both TimeScape and MapScape support the following mouseover functionality: branch mouseover highlights the descendant lineages; clone mouseover brings attention to the clone throughout the view and displays its prevalences at each time point or anatomical location; and phylogenetic class mouseover highlights the relevant tumour samples. The legend phylogeny supports clicking functionality: clicking a clone filters the mutation table for mutations originating in the selected clone. Note that the visualization supports the clicking of multiple clones, allowing the user to highlight specific combinations of clones throughout the view. Clicking the Reset button on the top panel exits a clonal selection.

The TimeScape and MapScape mutation tables house the patient's deep sequencing mutations. The table's Search box includes the capability to filter mutations by chromosome, coordinate, gene, etc. Clicking on a mutation highlights the legend phylogeny branch in which it occurred, and displays its variant allele frequency in each sample. Clicking the Reset button exits a mutation selection.

The main CellScape view shows a phylogeny and a heatmap. The top bar houses buttons with the following capabilities: toggling between the traditional and force-directed graph views; incorporating edge distances into the phylogeny; and selecting genomic profiles of interest to view the corresponding single cells in the phylogeny (note the support for multiple selections). The CellScape legend contains the following mouseover functionality: mouseover of an annotation class will display its associated single cells in the phylogeny and heatmap. When time-series information is provided as input to CellScape, a TimeScape will be appended to the view. The CellScape and TimeScape are interactively linked by mouseover of any clonal or temporal view elements.

#### Colouring the clonal phylogeny

Although the user may specify custom clone colours, the default colours are computed in a way that visually separates clonal lineages by hue. The algorithm, summarized by Figure S9, proceeds by first finding the *marker clones,* where a marker clone is a clone that marks the start of a lineage. The marker clones are assigned (in depth-first search order) a hue from a split colour wheel. All other clones obtain a darker version of their ancestor. The choice to darken rather than lighten a descendant clone reflects the accumulation, rather than the loss, of mutational content as lineages evolve.

#### Representing small clonal populations in TimeScape

Figure S10 shows that at any time point in TimeScape, clones with less than 0.5% prevalence are artificially vertically expanded to 0.5% of the view height, and the remaining clones experience a proportionate reduction in height. This ensures the visibility of low-prevalence clones, as well as the ability to discriminate between clonal minority and absence.

#### X-axis spacing of emergent clones in TimeScape

Prior to each time point in TimeScape, certain clones may emerge. The procedure for spacing these emergent clones along the x-axis is summarized by Figure S11. The height of the longest emergent lineage is used to evenly horizontally space the emergent clones in the previous window of time. If a perturbation event is specified between two time points, the spacing of emergent clones will be achieved after the event but before the next time point.

#### Vertical genotype layouts in TimeScape

Three genotype layouts—*stacked, centred* and *spaced*—are available as a user parameter. Figure S12 directly compares the three layouts. The default genotype layout in TimeScape is the stacked layout, which stacks genotypes one on top of the another (Fig. S12a, Supplementary Link 8). The spaced layout stacks the genotypes while ensuring a minimum amount of spacing such that each genotype is vertically surrounded by its ancestor (Fig. S12b, Supplementary Link 9). The centered layout centers genotypes with respect to their ancestors (Fig. S12b, Supplementary Link 10).

#### Perturbation events in TimeScape

To incorporate a perturbation event (Fig. 3a “Chemotherapy” time point) into TimeScape, the user specifies the event name, the subsequent time point, and the remaining fraction of total tumour content. During the event, the plot height will shrink to the specified fraction.

#### Auto-layout of samples in MapScape

Although the user may specify a custom sample order for the radial layout, the default sample order is dependent on the relationship between the sample's anatomic location and the centre of the main view—the line segment between these two points forms an angle with the x-axis, and this angle is used to order the samples in a clockwise manner starting at the positive x-axis (Fig. S13).

#### Image auto-crop in MapScape

The user-provided image is cropped automatically based on the user-specified sample coordinates. The auto-crop function ensures a minimum space of 15 pixels between the edge of the circular cut-out and the sample-associated anatomic locations.

#### Handling minor clones in MapScape

The statistical processing of clonal evolution datasets often assigns extremely small but non-zero clonal prevalences when in reality the clones are absent at particular anatomical sites. To display these minor clones would unnecessarily complicate the view. By default, therefore, clones with less than 1% prevalence are not shown, but warnings will be issued in the console, viewable by opening the browser inspector.

The user may wish to display minor clones by setting the “show_low_prev_gtypes” parameter to “TRUE”. In this case, the clones, where minor, will appear as open circles in the site-specific phylogeny. Note that a clone will only appear in the cellular aggregate if its prevalence is more than 1/“n_cells”, where “n_cells” is a parameter specifying the total number of cells to display in the aggregate.

#### Cellular aggregate layout in MapScape

The visualization of each sample contains a cellular aggregate representation of the tumour. Cellular aggregates are created by Voronoi tessellation, a mathematical process that partitions a plane into polygons such that each polygon contains one coordinate from a predetermined set, and the space within each polygon is closer to its containing coordinate than to any other coordinate within the set. The polygons, or “cells”, are assigned clonal colours proportionately to the clonal prevalences within the tumour. For a given patient, this colouration adheres to two principles: *stability,* where spatial regions are preserved (e.g. the green clone will always appear on the left-hand side of an aggregate), and *contiguity,* where each clone has a maximal clustering coefficient. Each clone is assigned a *starting cell* where their colouration will begin. These starting cells are evenly spaced along the aggregate's edge, and remain consistent across aggregates to ensure *stability.* To ensure *contiguity,* the assignment of colours to cells proceeds in a round-robin fashion, whereby each clone, in turn, is assigned to the uncoloured cell nearest to its starting cell.

#### Vertically ordering the single cells in the phylogeny and heatmap

In CellScape, the vertical plane is divided equally such that each observed cell occupies a unique vertical position. The vertical ordering of cells proceeds according to the phylogeny. Sibling cells are vertically ordered based on the size of their descendant lineage, and ancestors are centered vertically with respect to their descendants. Some of these ancestors are indicated by an empty circle—these are *latent nodes*, or inferred nodes that are not sampled. Any leaf nodes or peripheral lineages void of genomic information are removed from the phylogeny—in this situation, console warnings will indicate node removal.

#### Targeted mutation ordering in CellScape

Targeted mutations may be ordered manually by the user. The default, however, is to order them by the clonal phylogeny (if present), using a method summarized in Figure S14. In this method, each mutation is assigned a set of clones. A clone is assigned to a mutation if at least a parameterized proportion (default is 0.2) of single cells within the clone contain the mutation at a Variant Allele Frequency (VAF) above a parameterized threshold (default is 0.05). The mutations are then ordered by their clonal sets, which are in turn ordered by all combinations of a depth-first search of the clonal phylogeny (e.g. {A, B, C, D, E}). Within each clonal set, the mutations are sorted from left to right by their mean VAF for the entire dataset of single cells.

If neither a custom mutation order nor the clonal phylogeny are provided, the mutations are ordered by a hierarchical clustering method which clusters the mutation sites according to their VAFs for each cell (E.g. Fig. S15). Prior to clustering, VAFs are rounded: VAFs between 0.05 and 0.95 are given a heterozygous value of 0.5, and VAFs greater than or equal to 0.95 are given a homozygous value of 1. Also, VAFs less than 0.05 are defined as −10—upon clustering (using the “hclust” function in the “stats” R package, version 3.3.1), this separates mutation sites with few and many variants. Given the resultant mutation order, if the mean mutation VAF is increasing rather than decreasing, the order is reversed to ensure that ancestral mutations (high mean VAF) are on the left of the matrix, and newly acquired mutations (low mean VAF) are on the right of the matrix.

#### Sharing the visualizations

All E-scape visualizations may be exported as a static PNG or SVG by clicking the appropriate button on the top panel. In order to share the interactive visualizations directly, the user may call the TimeScape, MapScape or CellScape tools from within an RMark-down document, which will produce a shareable html document. Note that multiple interactive visualizations may be shared on the same html document.

## Acknowledgements

We wish to acknowledge the generous long term funding provided by the BC Cancer Foundation. The project was supported by a Canadian Cancer Society Research Institute Innovation grant and a Genome Canada/Genome BC Disruptive Innovation in Genomics grant to SPS. SPS is supported by Canada Research Chairs and is a Michael Smith Foundation for Health Research scholar. MAS and AM are supported by NSERC CGS scholarships. AR is supported by a CIHR CGS scholarship.

